# Prediction of survival after partial hepatectomy using a physiologically based pharmacokinetic model of indocyanine green liver function tests

**DOI:** 10.1101/2021.06.15.448411

**Authors:** Adrian Köller, Jan Grzegorzewski, Hans-Michael Tautenhahn, Matthias König

**Affiliations:** Institute for Theoretical Biology, Institute of Biology, Humboldt University, Berlin, Germany; Experimental Transplantation Surgery, Department of General, Visceral and Vascular Surgery, Jena University Hospital, Jena, Germany

**Author notes:** Correspondence: Matthias König.

**Keywords:** Indocyanine Green, Hepatectomy, Liver Cirrhosis, Mathematical Model, Computational Model, Pharmacokinetics, Liver Function, Liver Resection

## Abstract

The evaluation of hepatic function and functional capacity of the liver are essential tasks in hepatology as well as in hepatobiliary surgery. Indocyanine green (ICG) is a widely applied test compound that is used in clinical routine to evaluate hepatic function. Important questions for the functional evaluation with ICG in the context of hepatectomy are how liver disease such as cirrhosis alters ICG elimination, and if postoperative survival can be predicted from preoperative ICG measurements. Within this work a physiologically based pharmacokinetic (PBPK) model of ICG was developed and applied to the prediction of the effects of a liver resection under various degrees of cirrhosis. For the parametrization of the computational model and validation of model predictions a database of ICG pharmacokinetic data was established. The model was applied (i) to study the effect of liver cirrhosis and liver resection on ICG pharmacokinetics; and (ii) to evaluate the model-based prediction of postoperative ICG-R15 as a measure for postoperative outcome. Key results are the accurate prediction of changes in ICG pharmacokinetics caused by liver cirrhosis and postoperative changes of ICG-elimination after liver resection, as validated with a wide range of data sets. Based on the PBPK model, individual survival after liver resection could be classified, demonstrating its potential value as a clinical tool.

## 1 INTRODUCTION

Determining liver function is a crucial task in hepatology, e.g., for liver disease diagnosis or evaluation of pre- and postoperative functional capacity of the liver. An accurate assessment is especially relevant in the context of liver surgery as postoperative complications are associated with reduced functional capacity. A comprehensive characterization of the status of a patient and their liver is routinely performed before liver surgery such as partial hepatectomy. This includes among others anthropometric factors (e.g. age, sex, body weight), static liver function tests (e.g., ALT, AST, albumin, bilirubin, INR, prothrombin time), cardiovascular parameters (e.g. cardiac output, blood pressure, hepatic blood flow) and lifestyle factors (e.g. smoking, medication) as well as volumetric information obtained by CT or MRI scans. In addition, quantitative evaluation of liver function is often performed via pharmacokinetic measurements of test compounds specifically metabolized by the liver (dynamical liver function tests) such as methacetin (LiMAx (Rubin et al., 2017)) and MBT (Gorowska-Kowolik et al., 2017)), caffeine (Renner et al., 1984), galactose (Bernstein et al., 1960) and indocyanine green (ICG) (Sakka, 2018).

ICG is an inert, anionic, water-soluble, tricarbocyanine dye that is bound to plasma proteins. After intravenous administration ICG is taken up exclusively by the liver and excreted unchanged into the bile. It is not reabsorbed by the intestine and does not undergo enterohepatic circulation (Wheeler et al., 1958). As a result, ICG is an ideal test compound to test hepatic uptake and biliary excretion. Based on the plasma time course of ICG, pharmacokinetic parameters are calculated as a proxy for liver function, the most common parameters being: (i) ICG retention ratio 15 minutes after administration (ICG-R15) [%]; (ii) ICG plasma disappearance rate (ICG-PDR) [%/min]; (iii) ICG-clearance [ml/min]; and (iv) ICG half-life (ICG-t_1/2_) [min]. Reduced elimination of ICG by the liver is directly reflected by these parameters (Sakka, 2018).

Liver disease, especially advanced and more severe liver disease, is accompanied by a loss of liver function which can be quantified with dynamical liver function tests. The effects of liver disease on ICG-elimination have been studied extensively, e.g., in different stages of primary biliary cholangitis (PBC) (Vaubourdolle et al., 1991). ICG elimination is reduced in Gilbert’s disease (Martin et al., 1976) as well as in patients with hepatic fibrosis and cirrhosis (Gadano et al., 1997). Interestingly, also non-liver diseases can affect ICG parameters, e.g., ICG-clearance is significantly reduced in patients with chronic pancreatitis (Andersen et al., 1999).

Liver cirrhosis is an end stage liver disease and highly relevant in the context of hepatobiliary surgery. The most common causes for liver cirrhosis in Europe are alcohol abuse, chronic hepatitis B and/or C virus infection or non-alcoholic fatty liver disease (Hackl et al., 2016). The pathological characteristics of liver cirrhosis include degeneration of hepatocytes as well as a reduction of liver perfusion through increased portal resistance. An additional characteristics is the formation of intrahepatic shunts, which bypass part of the hepatic blood supply around the liver tissue. Consequently, no ICG can be extracted, resulting in further reduced elimination (Schuppan and Afdhal, 2008).

The severity of cirrhosis can be described using the Child-Turcotte-Pugh-Score (CTP) (Child and Turcotte, 1964; Pugh et al., 1973). The CTP is an empiric, qualitative, dis-continuous classification of the severity of the ”hepatic functional reserve” (Botero and Lucey, 2003). Based on a set of parameters, the CTP assigns a score from 5-15 to a cirrhotic patient, where the more severe a patients symptoms are the higher the score is. These parameters are serum bilirubin and serum albumin concentrations, the International Normalized Ratio (INR), amount of ascites in ultrasound and the degree of encephalopathy (Child and Turcotte, 1964; Pugh et al., 1973). Patients are classified depending on their mortality risk as CTP-A (5-6 points, low risk), CTP-B (7-9 points, intermediate risk) or CTP-C (10-15 points, high risk). Differences in ICG-elimination between cirrhotic patients and control subjects has been widely assessed (Caesar et al., 1961; Burns et al., 1991; Gilmore et al., 1982; Figg et al., 1995; Møller et al., 2019; Mukherjee et al., 2006; Pind et al., 2016). A good correlation between CTP-score and ICG-elimination has been reported with ICG-elimination decreasing as CTP-score increases (Figg et al., 1995; Møller et al., 2019; Mukherjee et al., 2006; Pind et al., 2016).

For many primary and secondary liver tumors, partial hepatectomy is the only curative treatment option. Liver resection (partial hepatectomy) describes the surgical removal of a part of the liver. Hepatectomy is an important procedure in general surgery with more than 20,000 liver resections in Germany per year (Filmann et al., 2019). The procedure has been widely performed for the treatment of various liver diseases, such as malignant tumors, benign tumors, calculi in the intrahepatic ducts, hydatid disease, and abscesses (Jin et al., 2013). Despite advances in technology and increasing numbers of highly experienced and specialized clinicians in the field of hepatobiliary surgery, postoperative morbidity and mortality is still a major issue as complex resections are increasingly being performed in older and higher risk patients (Jin et al., 2013). Major hepatectomy in the presence of cirrhosis is considered to be contraindicated due to the high mortality rate. Only selected patients with stable Child A status or an ICG-R15 of less than 10% may potentially be considered for extended liver resection. (Kitano and Kim, 1997).

A key challenge in HPB (hepato pancreato biliary) surgery is to predict functional capacity of the future liver remnant and thus reduce morbidity and mortality after extended liver resection. As a result, the decision whether or not a partial hepatectomy can be performed is often based on predictions of postoperative liver function (in addition to the remnant liver volume), which are in turn based on preoperative assessment of liver function (and volume). Understanding how cirrhosis alters liver function as measured via ICG is of high clinical relevance. Elucidating how ICG parameters change with increasing CTP score would be a valuable asset for the functional evaluation of patients with liver disease.

Important questions for the evaluation of liver function with ICG in the context of HPB surgery are how liver disease, especially cirrhosis, alter ICG elimination, and (ii) if postoperative survival can be predicted from preoperative ICG measurements. Within this work a physiologically based computational model of ICG pharmacokinetics was developed and applied to study these questions.

## 2 MATERIAL AND METHODS

### 2.1 Data

For the calibration and validation of the model a large data set of ICG measurements and physiological data was established. All data is available via the pharmacokinetics database PK-DB (https://pk-db.com) (Grzegorzewski et al., 2021). PK-DB was used to encode the information on (i) patient characteristics (e.g. age, disease, medication), (ii) applied interventions (e.g. ICG dosing, route of application); (iii) measured ICG time courses in plasma and (iv) ICG pharmacokinetic parameters (e.g. ICG-PDR, ICG-R15, ICG-clearance).

### 2.2 Indocyanine Green Pharmacokinetic Parameters

Pharmacokinetic parameters of ICG were calculated from the plasma-concentration time courses using non-compartmental methods (Urso et al., 2002). The elimination rate constant (*k*_*el*_) was calculated by fitting the concentration-decay-curve to an exponential function: 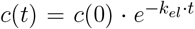. ICG-PDR is *k*_*el*_ reported in [%/min]. Half-life (*t*_1/2_) was calculated as *log*(2)*/k*_*el*_, ICG-clearance as *CL* = *V*_*d*_ *· k*_*el*_, with the apparent volume of distribution *V*_*d*_ = *D/*(*AUC*_*∞*_ = *kel*). *D* is the applied dose of ICG and *AUC*_*∞*_ is the area under the plasma-concentration time curve *AUC* calculated via the trapezoidal rule, extrapolated until infinity. ICG-R15 = *c*(15)*/c*_*max*_ was calculated as the ratio between the plasma-concentration after 15 minutes and the maximum concentration *c*_*max*_.

### 2.3 Model

The computational model is an ordinary differential equation (ODE) model encoded in the Systems Biology Markup Language (SBML) (Hucka et al., 2019; Keating et al., 2020). It is defined as a set of species (metabolites), compartments (organs and blood compartments) and reactions (processes such as metabolic reactions and blood transport). The model was developed using sbmlutils (Kö nig, 2021b), and cy3sbml (König and Rodriguez, 2019) and simulated using sbmlsim (König, 2021a) based on the high-performance SBML simulator libroadrunner (Somogyi et al., 2015).

### 2.4 Model Parameterization

Values for organ volumes and tissue blood flows were taken from literature (ICRP, 2002). A subset of model-parameters was determined by minimizing the residuals between model predictions and clinical data. This optimization-problem was solved using SciPy’s least squares method and differential evolution algorithm (Virtanen et al., 2020). For the objective cost function *F* depending on the parameters 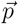 a simple L2-Norm was used consisting of the sum of weighted residuals

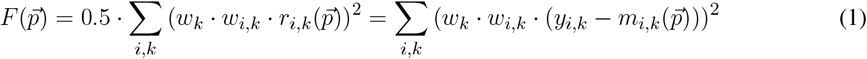

where 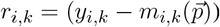 is the residual of time point *i* in time course *k* for model prediction 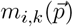 and the corresponding data point *y*_*i,k*_; *w*_*i,k*_ is the weighting of the respective data point *i* in time course *k* based on the error of the data point and *w*_*k*_ = the weighting factor of time course *k*. Weighting of time courses was based on the number of subjects per study. The final parameter set given in Tab. 2 was determined using 250 runs of the local least square optimization. The data used for the parameter fit is listed in Tab. 1.

**Table 1.** Overview of curated clinical studies.

| Study | PK-DB | PMID | Protocol | Body weight | Fit | Data used in fit | Description |
| --- | --- | --- | --- | --- | --- | --- | --- |
| Andersen1999 | PKDB00386 | 10499483 | Bolus: 0.5 mg/kg | Contr.: 83.4 kg;<br>Panc. 58.2 kg | ✓ | ICG time course (plasma) | Pharmacokinetic time course of ICG-disappearance in chronic pancreatitis and healthy subjects. |
| Burns1991 | PKDB00388 | 1848168 | Bolus: 0.5 mg/kg; Infusion: 0.25 mg/min | NR | ✓ | ICG time course (plasma) | ICG pharmacokinetics in healthy subjects and patients with liver disease. |
| Caesar1961 | PKDB00389 | 13689739 | Bolus: 0.5 mg/kg; Infusion: 0.5 mg/min | NR | ✓ | ICG extraction-ratio | Measuring hepatic blood flow and assessing hepatic function by ICG pharmacokinetics. |
| Cherrik1960 | PKDB00390 | 13809697 | Bolus: 0.5 mg/kg | NR | - | - | ICG pharmacokinetics in healthy subjects and patients with liver disease. |
| Chijiwa2000 | PKDB00391 | 10773154 | Bolus: 0.5 mg/kg | NR | ✓ | ICG time course (bile excretion) | Biliary excretion of ICG and ATP-dependency of ICG pharmacokinetics. |
| Figg1995 | PKDB00393 | 8602375 | Bolus: 0.5 mg/kg | NR | - | - | Comparison of methods to assess hepatic function including CTP-score and ICG-clearance |
| Gadano1997 | PKDB00394 | 9083919 | Infusion: 0.4 mg/min (controls); 0.8 mg/min (cirrhotics) with priming dose (24 mg controls; 12 mg cirrhotics). | NR | ✓ | - | ICG-clearance and extraction-ratio and their dependency on hepatic bloodflow in healthy subjects and in patients of hepatic fibrosis and cirrhosis. |
| Grainger1983 | PKDB00395 | 6822056 | Bolus: 0.25 mg/kg | NR | ✓ | ICG extraction-ratio | Non-invasive measurement of hepatic blood flow using ICG extraction-ratio. |
| Grundmann1992 | PKDB00396 | 1482735 | Bolus: 0.3 mg/kg | NR | ✓ | ICG time course (plasma) | Effect of anesthetics on ICG-pharmacokinetics and hepatic blood flow. |
| Herold2001 | PKDB00397 | 11169069 | Bolus: 0.5 mg/kg | NR | - | - | Comparison of liver function tests. Correlation between CTP-score and ICG-kel in healthy subjects and cirrhotic patients. |
| Kamimori2000 | PKDB00299 | 10883415 | Bolus: 0.5 mg/kg | 58.6 ± 11.2 kg | ✓ | ICG time course (plasma) | Effect of the menstrual cycle on ICG pharmacokinetics. |
| Keiding1993 | PKDB00402 | 8151094 | Infusion: 0.08 mg/min | NR | - | - | Effect of changing plasma protein concentrations on ICG-extraction. |
| Klockowski1990 | PKDB00403 | 2146057 | Bolus: 0.5 mg/kg | NR | ✓ | ICG time course (plasma) | Effect of isradipine and diltiazem on ICG-pharmacokinetics. |
| Leevy1962 | PKDB00404 | 14463639 | Infusion: 0.3 and 1.5 mg/min/m<sup>2</sup> with priming dose (10 mg). | NR | ✓ | ICG extraction-ratio; ICG time course (hv and ar) | Estimation of hepatic blood flow using ICG. |
| Leevy1967 | PKDB00405 | 6071462 | Bolus: 0.5 mg/kg and 5.0 mg/kg | NR | - | - | Estimation of liver function with ICG. Dose dependency of ICG-PDR. |
| Martin1975 | PKDB00406 | 1208580 | Bolus: 0.5, 2.0, 3.5 and 5.0 mg/kg | NR | - | - | Differences in ICG-pharmacokinetics in men and women at varying ICG-doses. |
| Martin1976 | PKDB00407 | 814028 | Bolus: 0.5, 2.0, 3.5 and 5.0 mg/kg | NR | - | - | ICG-pharmacokinetics in patients with Gilbert's Syndrome. |
| Meijer1988 | PKDB00408 | 3181282 | Bolus: 0.5, 1.0 and 2.0 mg/kg | 64.8 kg | ✓ | ICG time course (plasma and bile excretion) | Biliary excretion of ICG at different doses. |
| Moeller1998 | PKDB00409 | 9691928 | Infusion: Rate not reported | 72.7 (40-115) kg | - | - | Arterial hypoxaemia in cirrhosis. Correlation between ICG-clearance and CTP-score. |
| Moeller2019 | PKDB00410 | 30221390 | Infusion: 0.2 mg/min with priming dose (2 mg), Bolus: 0.5 mg/kg | 79.2 ± 18.5 kg | - | - | Correlation between ICG-pharmacokinetics and CTP-score. |
| Niemann2000 | PKDB00414 | 10801242 | Bolus: 10 mg | 80 ± 17 kg | ✓ | ICG time course (plasma) | ICG- pharmacokinetic time course under administration of propranolol. |
| Ohwada2006 | PKDB00412 | 16498606 | Bolus: 20 mg | NR | - | - | Prediction of postoperative liver functional capacity based on ICG-pharmacokinetics. |
| Okochi2002 | PKDB00415 | 11855925 | Bolus: 20 mg | NR | - | - | Comparison of preoperative and postoperative ICG-pharmacokinetics. Prediction of postoperative liver functional capacity based on ICG-pharmacokinetics. |
| Seyama2009 | PKDB00416 | 19208031 | Bolus: 0.5 mg/kg | NR | - | - | Assessment of liver function for safe hepatic resection. Prediction of survival after hepatectomy based on ICG-R15. |
| Soons1991 | PKDB00411 | 1768562 | Infusion: 2.0, 0.5, 1.0 mg/min (consecutive) | 72 ± 6 kg | ✓ | ICG time course (plasma) | Assessment of hepatic blood flow in healthy subjects by continuous infusion of ICG. |
| Stockmann2009 | PKDB00417 | 19561474 | Bolus: 0.5 mg/kg | NR | - | - | Prediction of postoperative liver functional capacity based on ICG-pharmacokinetics. |
| Sunagawa2021 | PKDB00418 | 33052632 | Bolus: 20 mg | NR | - | - | Prediction of postoperative liver failure based on ICG-kel measurements. |
| Thomas2015 | PKDB00419 | 25581073 | Bolus: 0.25 mg/kg | NR | - | - | Intraoperative prediction of postoperative liver functional capacity using trial-clamping and ICG-pharmacokinetics. |
| Wakabayashi2004 | PKDB00420 | 15013363 | Bolus: 0.5 mg/kg | NR | - | - | Correlation between preoperative ICG-R15 and estimated liver remnant. Prediction of survival based on ICG-R15. |

**Table 2.** Overview of key model parameters.

| Parameter | Description | Value | Unit | Fitted |
| --- | --- | --- | --- | --- |
| *BW* | Body weight | 75 | kg | - |
| *COBW* | Cardiac output per body weight | 0.83 | ml/s/kg | - |
| *QC* | Cardiac output ($BW \cdot COBW$) | 3.75 | l/min | - |
| *HCT* | Hematocrit | 0.51 | - | - |
| *Fblood* | Fraction of organ volume that is blood vessels | 0.02 | - | - |
| *FVgi* | Fractional tissue volume gastrointestinal tract | 0.0171 | l/kg | - |
| *FVli* | Fractional tissue volume liver | 0.0210 | l/kg | - |
| *FVlu* | Fractional tissue volume lung | 0.0076 | l/kg | - |
| *FVve* | Fractional tissue volume venous blood | 0.0587 | l/kg | - |
| *FVar* | Fractional tissue volume arterial blood | 0.0184 | l/kg | - |
| *LI_ICGIM_Vmax* | $V_{max}$ of liver import | 3.70E-2 | mmole/min/l | ✓ |
| *LI_ICGIM_Km* | $K_m$ of liver import | 2.17E-2 | mM | ✓ |
| *LI_ICGLI2CA_Vmax* | $V_{max}$ of bile excretion | 9.44E-4 | mmole/min/l | ✓ |
| *LI_ICGLI2CA_Km* | $K_m$ of bile excretion | 1.24E-2 | mM | ✓ |
| *LI_ICGLI2BI_Vmax* | $V_{max}$ of bile transport | 1.14E-4 | l/min | ✓ |

### 2.5 Uncertainty analysis

To evaluate the uncertainty of model predictions uncertainty analysis was performed for a subset of simulations. Each model parameter was changed individually by ±25%. From the set of resulting time courses the mean, standard deviation (SD) and minimum and maximum values at each time point were calculated. These uncertainty areas were displayed as shaded areas. Parameters corresponding to physical constants (such as molecular weights) and dosing were not varied in the uncertainty analysis, as well as parameters for conservation conditions such as the fractional blood flow through the lung (must be 1).

### 2.6 Classification

Classification models were developed which predict survival after hepatectomy (binary classification: Survivors/Non-Survivors) based on different sets of features (see Tab. 3). Classification was performed using scikit-learn (Pedregosa et al., 2011) with a C-support vector classifier using a polynomial kernel. For model training and evaluation a dataset of 141 patients with information on survival status, resection rate and preoperative ICG-R15 was used (Seyama and Kokudo, 2009; Wakabayashi et al., 2004). Cross-validation was performed using the ShuffleSplit method with 200 iterations and a train-test-ratio of 75%/25%. Based on the confusion matrix the following evaluation metrics were calculated: precision/positive predictive value (PPV), recall, balanced accuracy, F1 score, negative predictive value (NPV) and receiver operator curves (ROC).

For the classification models PBPK1 and PBPK2 the model parameter *f*_*cirrhosis*_ was determined from preoperative ICG-R15 values using

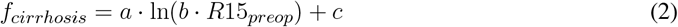

with *R*15_*preop*_ being the clinically measured preoperative ICG-R15 [dimensionless]. The coefficients *a* = 0.312, *b* = 1.693 and *c* = 0.861 were determined by fitting the curve to the predicted dependency of ICG-R15 on *f*_*cirrhosis*_ (see Fig. 4C). This provided an estimate of individual liver disease (cirrhosis degree). This estimated parameter allowed in combination with the resection rate to predict individual postoperative ICG-R15 values.

**Table 3.**
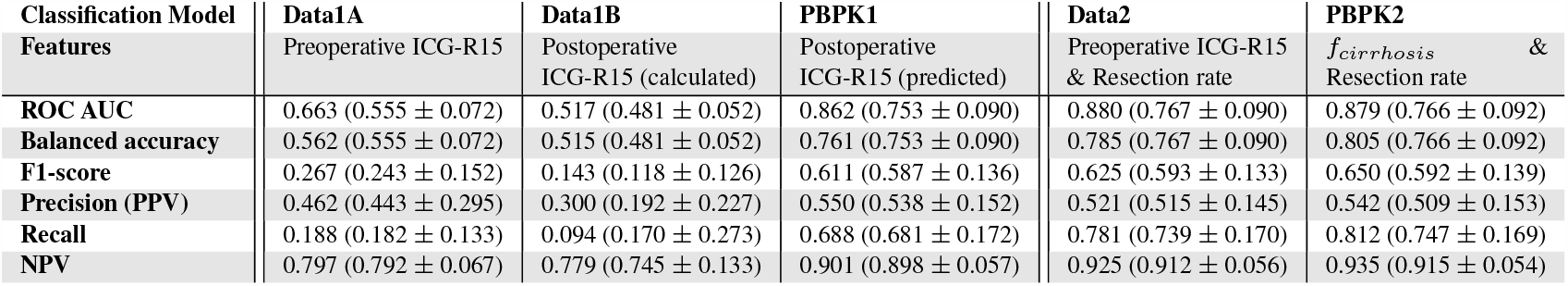
Evaluation metrics for classification models of survival after hepatectomy. Evaluation metrics of classification model are values for the model fitted with the complete dataset. Mean ± SD of cross validation reported in brackets.

## 3 RESULTS

Within this work a PBPK model of ICG pharmacokinetics was developed and applied to study (i) how liver disease, especially cirrhosis, alter ICG elimination, and (ii) if postoperative survival can be predicted from preoperative ICG measurements in the context of partial hepatectomy.

### 3.1 Data

A wide range of heterogeneous data was curated for model building (parameterization) and subsequent model validation (comparison of model predictions to clinical data). An overview of the 29 studies with their respective clinical protocols is provided in (Tab. 1). All data is freely available from https://pk-db.com.

### 3.2 Model

A PBPK model for the prediction of ICG pharmacokinetics was developed. To simulate the whole-body distribution and hepatic elimination of ICG two models were coupled: (i) A whole-body model (Fig. 1A) describing the distribution of ICG in the body and to the organs via blood flow. (ii) A liver model (Fig. 1B) which describes hepatic uptake of ICG, biliary excretion of ICG and transport of ICG into the feces. ICG pharmacokinetic parameters (i.e. ICG-PDR, R15, clearance, half-life) were calculated from the resulting time course predictions of ICG in the venous plasma.

**Figure 1.**
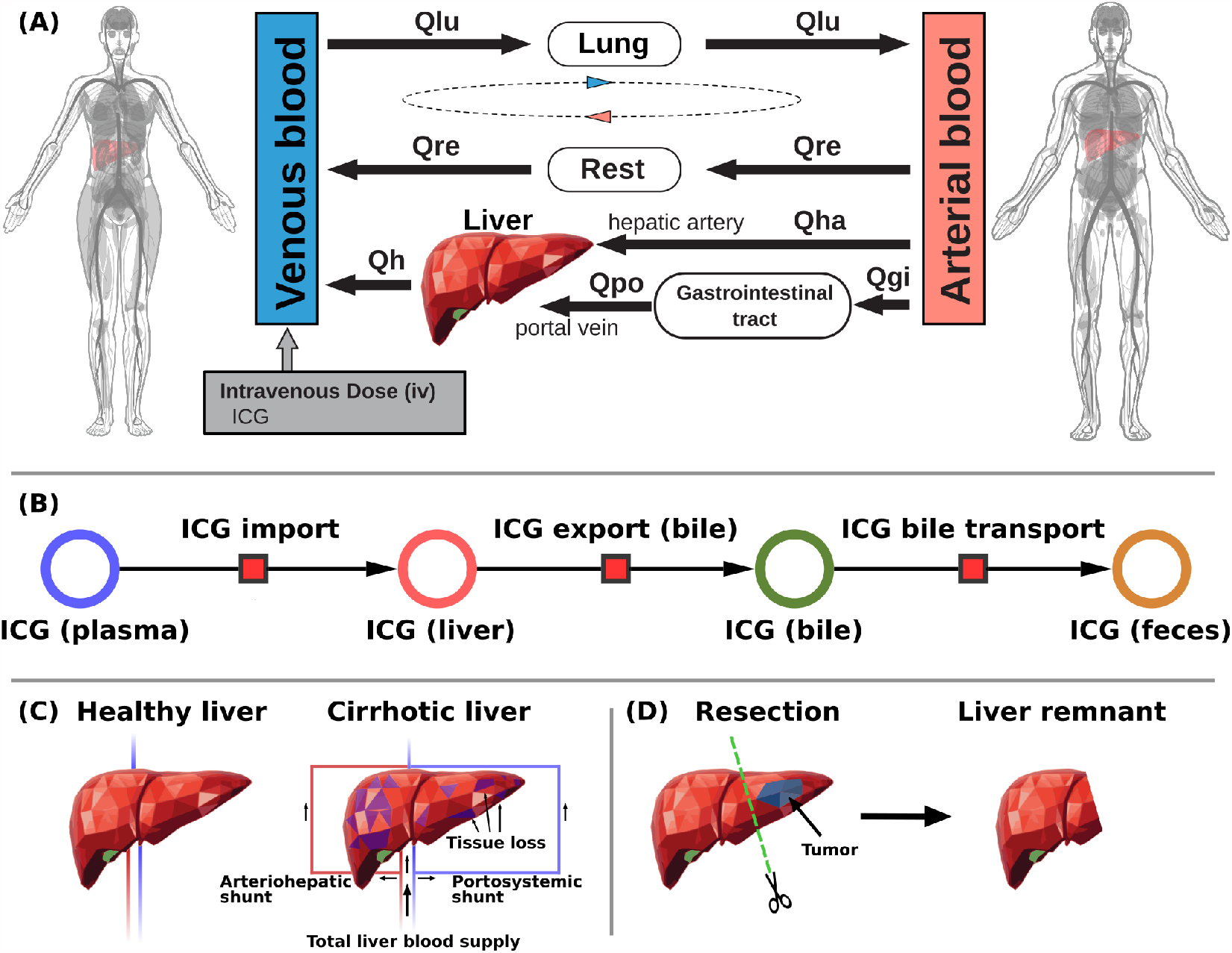
Model overview: **A:** Whole-body model. The whole-body PBPK model for ICG consists of venous blood, arterial blood, lung, liver, gastrointestinal tract and rest compartment (accounting for organs not modeled in detail) and the systemic blood circulation connecting these compartments. **B:** Liver model. ICG in the liver plasma compartment is taken up into the liver tissue (hepatocytes). Subsequently hepatic ICG is excreted in the bile from where it is excreted in the feces. No metabolization of ICG takes place in the liver. **C:** Modeling liver cirrhosis. Liver cirrhosis was modeled as a combination of tissue loss and hepatic shunts (see main text for details). **D:** Modeling hepatectomy. Hepatectomy was modeled as a removal of tissue volume with corresponding vessels (see main text for details)

#### 3.2.1 Distribution and blood flow

The distribution of ICG on the whole-body level is modeled using a network of blood flows representing the systemic circulation. From the venous blood, ICG is transported through the lung into the arterial blood from where it can reach the liver on two paths: (i) through the hepatic artery and (ii) through the gastrointestinal tract via the portal vein. Because the liver is the only tissue partaking in the elimination of ICG, all other organs (e.g. kidney, heart, adipose tissue, muscle, etc.) were pooled into the rest compartment. Each organ consists of a blood compartment (representing the vessels) and a tissue compartment. ICG transport via blood flow was implemented as irreversible transport. The transport *v*_*i*_ from compartment *i* to the next compartment is determined by the ICG concentration *C*_*i*_ in compartment *i* and a compartment-specific blood flow *Q*_*i*_. *Q*_*i*_ is determined by the cardiac output *Q*_*CO*_ and a compartment specific fractional tissue blood flow *f Q*_*i*_. Multiple conservation conditions hold in the model to ensure mass and flow balance. First the sum of blood flows from the arterial to the venous compartment must equal the sum of flows in the opposite direction: *Q*_*CO*_ = *Q*_*lu*_ = *Q*_*h*_ + *Q*_*re*_. Flow into an organ must be equal to the flow out of the organ. E.g. hepatic venous blood flow must be equal to the sum of hepatic arterial and portal venous blood flow: *Q*_*h*_ = *Q*_*ha*_ + *Q*_*po*_.

#### 3.2.2 Hepatic metabolism and biliary excretion

The liver model (Fig. 1B) consists of three consecutive transport reactions of ICG. After ICG is taken up in the liver it is excreted into the bile. Both transport reactions are modeled as irreversible Michaelis-Menten-kinetics. From the bile, ICG is transported into the feces modeled via a first order kinetic. All transport kinetics scale with the liver volume *V*_*li*_.

#### 3.2.3 Parameter fitting

Parameter fitting of the model was performed using a subset of ICG time courses and extraction-ratio measurements (see Tab. 1). No ICG pharmacokinetic parameters were used in the model fitting. Overall, 5 model parameters were fitted (see Tab. 2). Two of them determine the import of ICG in the liver, three determine the subsequent excretion in the bile. The agreement between fit data and model predictions improved substantially during parameter fitting and all training data with the exception of three simulations (Meijer1988, Chijiiwa2000 and Burns1991) could be described very well after parameter fitting.

#### 3.2.4 Modeling liver cirrhosis

The reference model, representing a healthy human subject, was adjusted to simulate cirrhosis by including a combination of functional tissue loss (due to scarring and necrosis in cirrhosis) and the formation of intrahepatic shunts, both key hallmarks of cirrhosis (Fig. 1C). The loss of functional liver tissue was controlled via the parameter *f*_*tissue_loss*_ ∈ [0, 1) which defines the fraction of parenchymal cell volume lost in the liver due to the disease. For modeling arteriohepatic and portosystemic shunts two additional blood vessels were introduced into the model. They connect the hepatic artery and the portal vein directly to the hepatic vein. As a result, a part of the portal venous and arterial blood bypasses the active liver tissue and is shunted to the hepatic venous blood compartment, so that ICG can not be extracted (corresponding to *in silico* shunts). The amount of blood that flows through the shunts is determined by the parameter *f*_*shunts*_ ∈ [0, 1), which defines the fraction of blood bypassing the liver. The remaining blood (1 − *f*_*shunts*_) reaches the liver tissue and ICG can be extracted. To simulate various degrees of cirrhosis the parameters *f*_*shunts*_ and *f*_*tissue_loss*_ were varied in lockstep by coupling them into the parameter *f*_*cirrhosis*_. The following values for *f*_*cirrhosis*_ were used: healthy - 0.0, mild cirrhosis - 0.41, moderate cirrhosis - 0.70, severe cirrhosis - 0.82.

#### 3.2.5 Modeling hepatectomy

The developed model allows to predict changes in ICG pharmacokinetic parameters after partial hepatectomy (Fig. 1D). *In silico* liver resections were simulated by reducing the fractional liver volume *FVli* by up to 90% (corresponding to a resection rate of 90%). The absolute liver volume is determined with the body weight *BW* via *FVli · BW*. All liver resections were simulated under varying degrees of cirrhosis as described above.

### 3.3 Healthy controls

In a first step the fitted model was evaluated with the data used for model calibration consisting of ICG time courses in healthy subjects (Fig. 2). For the simulations infusion protocols and body weights were adjusted as reported in the respective studies (see Tab. 1 for details). If no body weight was reported 75 kg were assumed.

**Figure 2.**
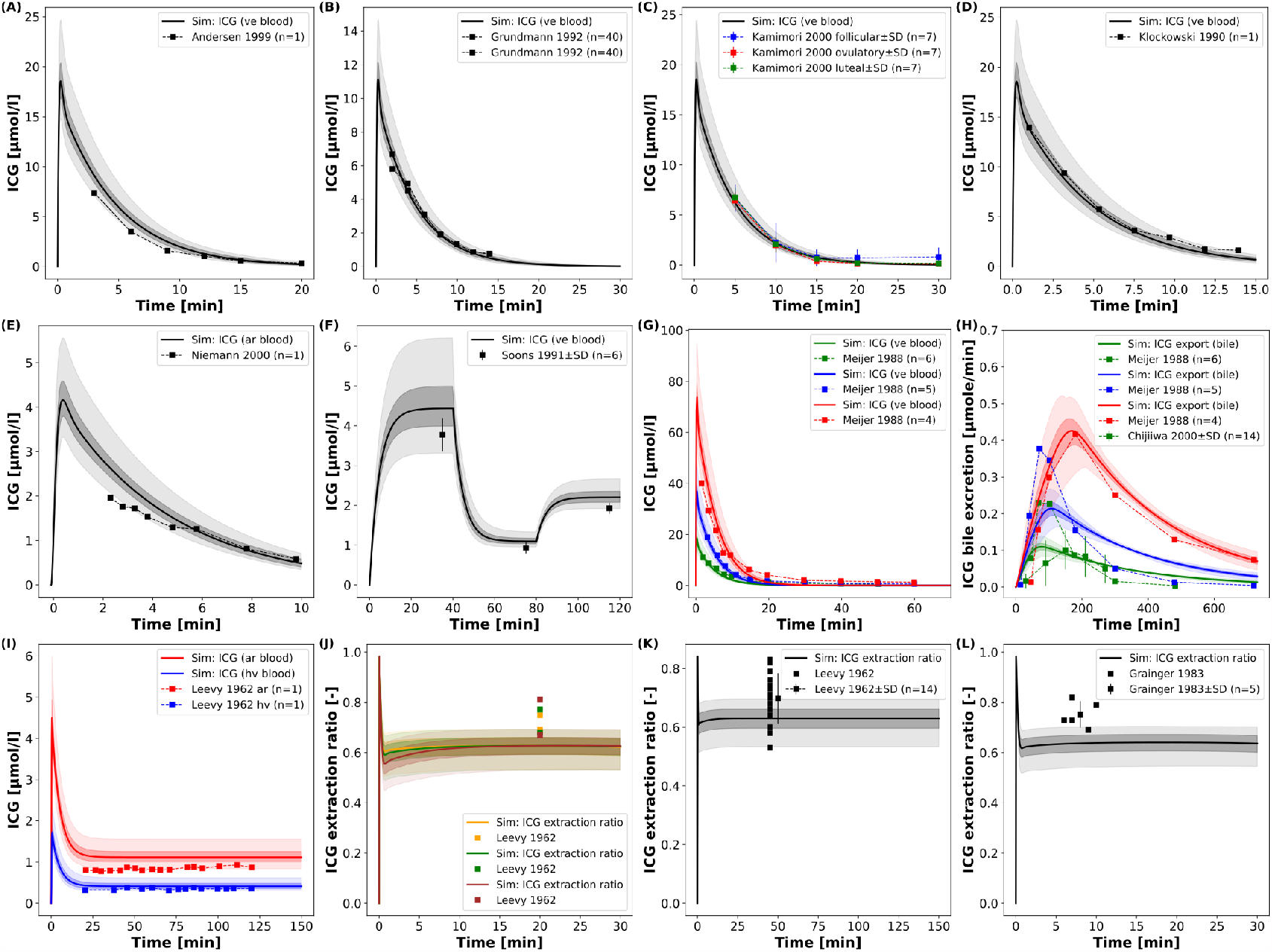
Model prediction of ICG time courses in healthy subjects: **A-D:** Venous concentration after bolus ICG administration (Andersen et al., 1999; Grundmann et al., 1992; Kamimori et al., 2000; Klockowski et al., 1990). **E:** Arterial concentration after bolus ICG administration (Niemann et al., 2000). **F:** Venous concentration during an ICG infusion protocol (2.0, 0.5, 1.0 mg/min, 40 minutes each) (Soons et al., 1991). **G, H:** Venous concentration and biliary excretion rate after 3 different ICG doses (0.5, 1.0, 2.0 mg/kg) (Meijer et al., 1988; Chijiiwa et al., 2000). **I:** Hepatic venous and arterial concentration during constant ICG infusion (Leevy et al., 1962). **J-L:** ICG extraction ratio during constant infusion (Leevy et al., 1962; Grainger et al., 1983).

The model predictions for ICG plasma disappearance curves after an ICG bolus are in good agreement with the clinical data (Andersen et al., 1999; Grundmann et al., 1992; Kamimori et al., 2000; Klockowski et al., 1990; Niemann et al., 2000; Meijer et al., 1988) (Fig. 2A-E). In addition, more complex infusion protocols as reported in Soons et al. (Soons et al., 1991) can also be described (Fig. 2F), infusion protocol of three different infusion rates (2.0 → 0.5 → 1.0 mg/min, each for 40 minutes). Due to the high extraction-ratio of ICG by the liver, the plasma concentration reaches steady state quickly after each change in the infusion rate. Next, simulations of the biliary excretion rate of ICG after bolus administrations of 0.5, 1.0 and 2.0 mg/kg ICG were performed and the results were compared to clinical data (Meijer et al., 1988; Chijiiwa et al., 2000) (Fig. 2G, H).

Finally, simulations of constant infusions were performed and compared to reported arterial and hepatic vein time courses of ICG (Leevy et al., 1962) and ICG extraction ratios (Leevy et al., 1962; Grainger et al., 1983) (Fig. 2I-L).

Overall, the model shows the ability to accurately predict ICG time courses for venous and arterial plasma concentrations, for hepatic vein concentrations, the biliary excretion rate and extraction ratios when compared to clinical data. Especially plasma time courses of ICG after ICG bolus and ICG infusion are very well predicted by the model, even for varying administration protocols (dosing and infusion rates).

In a next step a systematic analysis of the dose dependency of ICG pharmacokinetic parameters was performed (Fig. 3).A dose-dependency of the ICG parameters can only be observed if the ICG dose exceeds 100 mg (much higher then the typically applied doses of 20 - 35 mg), resulting in a reduction in ICG-clearance and ICG-PDR as well as an increase of ICG-R15 and ICG-t_1/2_ (Fig. 3A-D). The model predictions could be validated with clinical data (Martin et al., 1975, 1976; Meijer et al., 1988) (Fig. 3E-G).

**Figure 3.**
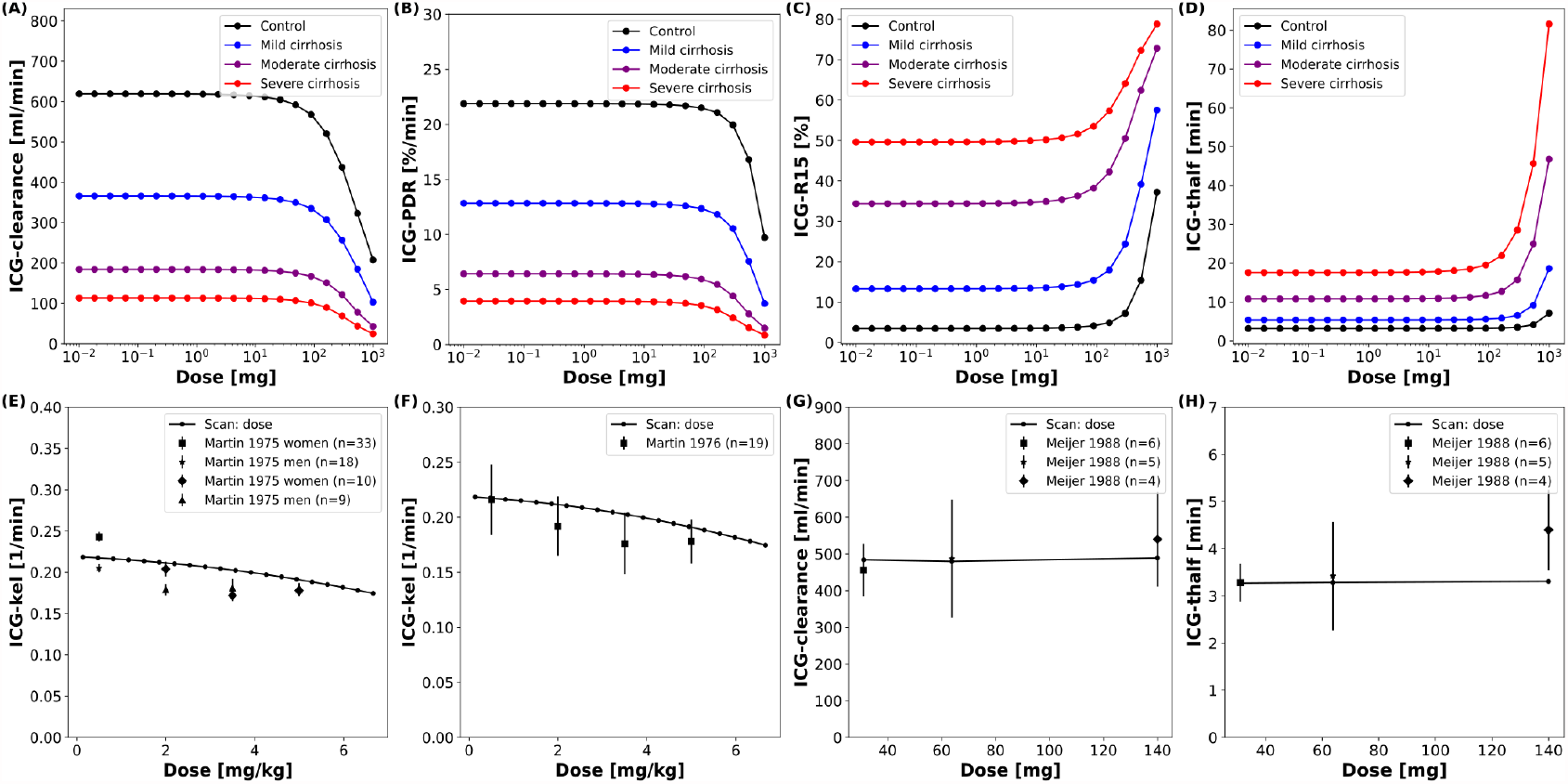
Dose dependency of ICG pharmacokinetic parameters: **A-D:** Dose dependency of ICG pharmacokinetic parameters in controls and three different degrees of cirrhosis. **E-H:** Dose dependency of ICG-kel, ICG-clearance, ICG-t_1/2_ in healthy subjects with clinical data (Martin et al., 1975, 1976; Meijer et al., 1988).

### 3.4 Cirrhosis

To simulate changes of ICG pharmacokinetics in cirrhosis, hepatic tissue loss and shunts were included in the model as described above. First, a systematic analysis of the effect of intrahepatic shunts (*f*_*shunts*_), functional tissue loss (*f*_*tissue_loss*_) and the combination of both (*f*_*cirrhosis*_) on ICG pharmacokinetic parameters was performed (Fig. 4A-D). All three parameters were varied from 0 (no effect, healthy control) to 0.9 (severe effect). ICG-clearance and ICG-PDR decrease with increasing *f*_*cirrhosis*_ whereas ICG-R15 and ICG-t_1/2_ increase. The loss of a fraction of functional liver tissue appears to have a smaller effect on ICG pharmacokinetic parameters than shunting of an equal fraction of blood past the liver. When *f*_*shunts*_ and *f*_*tissue_loss*_ are combined to *f*_*cirrhosis*_ their effect on ICG pharmacokinetic parameters is additive. For ICG-clearance and ICG-PDR the effects of both parameters combine to an almost linear dependency on *f*_*cirrhosis*_. The decrease in ICG-clearance and ICG-PDR with increasing cirrhosis and increase of ICG-R15 and ICG-t_1/2_ with increasing cirrhosis can be observed over a wide range of applied ICG doses (Fig. 3A-D).

**Figure 4.**
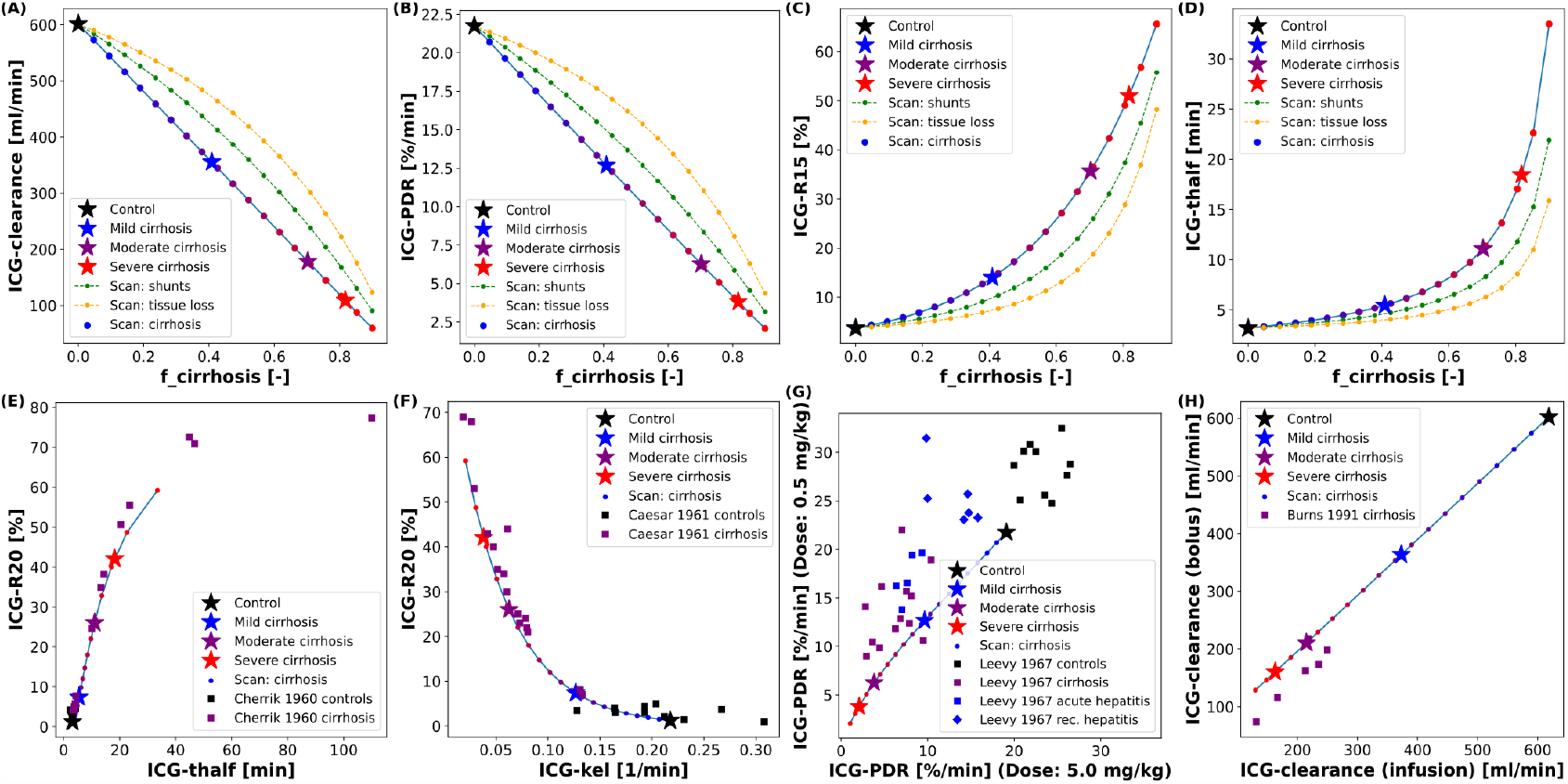
Dependency of ICG pharmacokinetic parameters on cirrhosis.: Simulations of 4 specific cirrhosis degrees are indicated by stars. **A-D:** Dependency of ICG pharmacokinetic parameters on the degree of shunting (green), degree of tissue loss (yellow) and degree of cirrhosis (black-blue-red). **E:** Correlation between ICG-R20 and ICG-t_1/2_ in cirrhotic and control subjects (Cherrick et al., 1960). **F:** Correlation between ICG-R20 and ICG-kel in cirrhotic and control subjects (Caesar et al., 1961). **G:** Correlation between ICG-PDR after an ICG dose of 0.5 mg/kg and 5.0 mg/kg in control subjects and subjects with various liver diseases (Leevy et al., 1967). **H:** Correlation between ICG-clearance after a bolus administration and during a constant infusion of ICG in cirrhotic subjects (Burns et al., 1991).

By varying the *f*_*cirrhosis*_ parameter from 0 to 0.9 different degrees of cirrhosis were simulated and the nonlinear relation between ICG-R20 and ICG-kel as well as ICG-R20 and ICG-t_1/2_ could be predicted (Fig. 4E,F). As seen in the systematic analysis (Fig. 4A-D) ICG-t_1/2_ and ICG-R20 increase with cirrhosis whereas ICG-kel decreases. The correlation between the ICG pharmacokinetic parameters is predicted accurately by the model when compared to a clinical dataset that lacks information about the severity of liver cirrhosis of its patients(Cherrick et al., 1960; Caesar et al., 1961). Next the ICG-PDR in cirrhotic patients, acute and recovering hepatitis and control subjects after different doses of ICG (0.5 mg/kg and 5.0 mg/kg ICG) was compared to the model predictions. The clinical data shows higher ICG-PDR values after an ICG dose of 0.5 mg/kg than after an ICG dose of 5.0 mg/kg (Leevy et al., 1967). In the model prediction the ICG-PDR in acute and recovering hepatitis resembles that of mild to moderate cirrhosis (Fig. 4G). ICG-clearances after a bolus administration and during a constant infusion show good positive correlation in cirrhotic patients (Burns et al., 1991). This correlation is predicted accurately by the model (Fig. 4H).

Having evaluated and validated the effect of *f*_*cirrhosis*_ on the model prediction of ICG parameters, we were interested how the model *f*_*cirrhosis*_ parameter compares to the *in vivo* estimation of cirrhosis degree via the CTP-score (Fig. 5). As described above, the CTP-score is a semi-quantitative scoring system that describes the severity of liver cirrhosis. An important step to apply the developed PBPK model in a clinical setting, is the ability to adjust the model individually to the respective status of liver disease in a patient. Therefore, the relationship between the *f*_*cirrhosis*_ parameter and the CTP-Score was evaluated using multiple datasets in which ICG pharmacokinetic parameters were reported in patient subgroups of different CTP-Scores (Figg et al., 1995; Møller et al., 1998, 2019; Herold et al., 2001).

**Figure 5.**
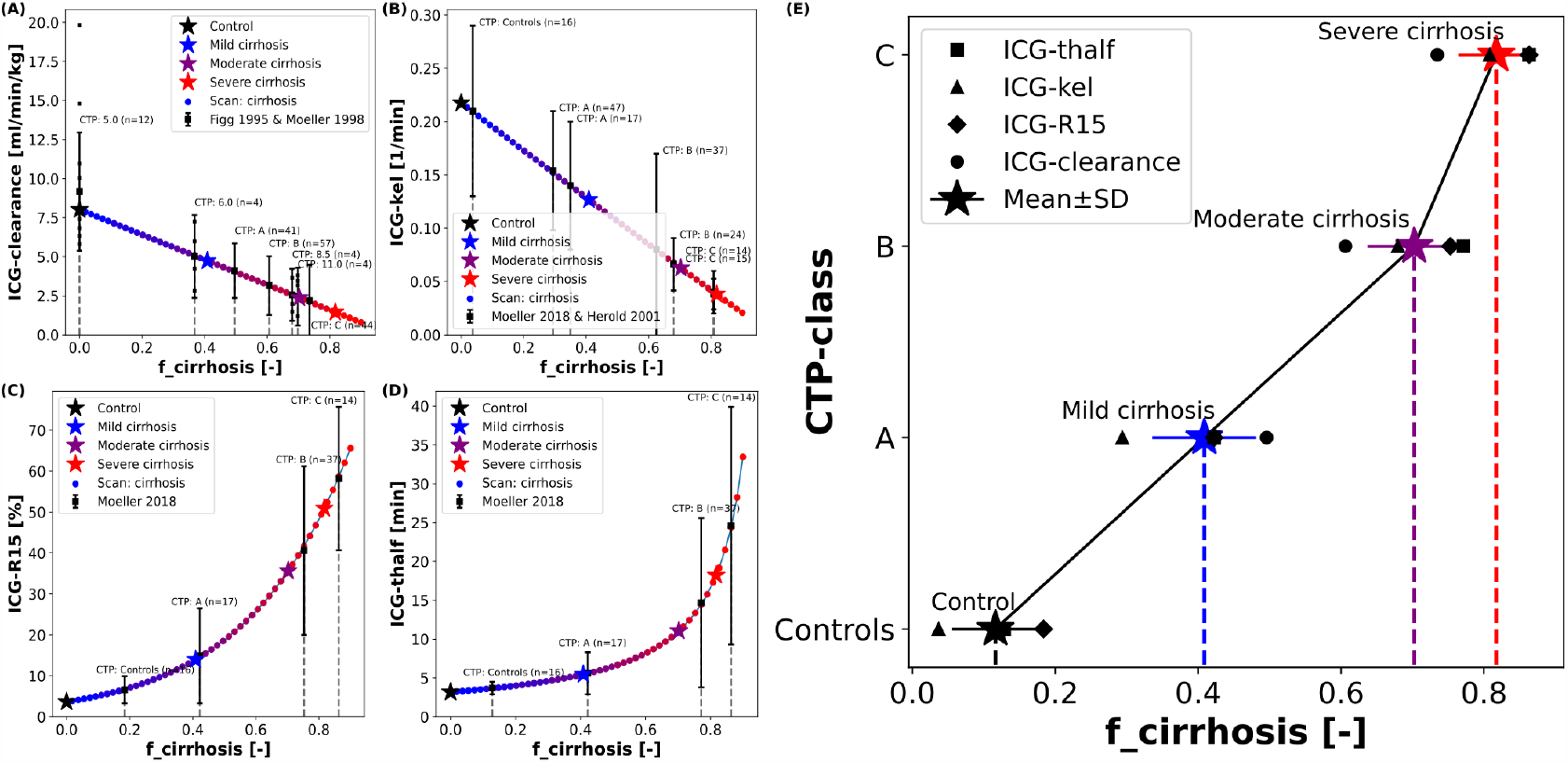
Mapping of model cirrhosis degree on CTP-score: **A:** Mapping based on ICG clearance (Figg et al., 1995; Møller et al., 1998). **B:** Mapping based on ICG-kel (Møller et al., 2019; Herold et al., 2001). **C:** Mapping based on ICG-R15 (Møller et al., 2019). **D:** Mapping based on ICG-thalf (Møller et al., 2019). **E:** Resulting *f*_*cirrhosis*_ values for each CTP-class combining the information from the mappings based on individual ICG pharmacokinetic parmeters (Control: *f*_*cirrhosis*_ = 0.0; Mild cirrhosis: 0.41; Moderate cirrhosis: 0.70; Severe cirrhosis: 0.82).

The clinical results of the ICG pharmacokinetic parameters in different CTP-classes were mapped onto their respective systematic scan (Fig. 5A-D). The resulting *f*_*cirrhosis*_ values were then compared between the patient groups. Additional individual data is shown (Figg et al., 1995).

The resulting mapping between *f*_*cirrhosis*_ and the CTP-classes shows a good positive correlation (Fig. 5E). The *f*_*cirrhosis*_ values for the controls groups are close to 0, increasing with the CTP-class. The relation appears nonlinear, as *f*_*cirrhosis*_ shows little difference between CTP-class B and C. The mappings of the CTP-class to *f*_*cirrhosis*_ for the different ICG parameters each give very similar results. From the mapping of all four pharmacokinetic parameters a mean value of *f*_*cirrhosis*_ was calculated for each CTP-class (Control: *f*_*cirrhosis*_ = 0.0; Mild cirrhosis: 0.41; Moderate cirrhosis: 0.70; Severe cirrhosis: 0.82). The resulting values were used in all simulations of control, mild, moderate and severe cirrhosis, as well as in the above described dose dependency analysis (Fig. 3A-D).

Only a single study reported the numerical CTP-score of the patient groups in combination with ICG-clearance (Figg et al., 1995). All other studies instead used the CTP-classes (A, B, C) (Møller et al., 1998, 2019; Herold et al., 2001). With a dataset of individually reported CTP-scores in combination with ICG pharmacokinetic parameters of cirrhotic patients, it would be possible to calculate the relationship of the CTP-score on the *f*_*cirrhosis*_ parameter more accurately. Such an improved mapping would allow to adjust the model via the *f*_*cirrhosis*_ parameter individually based on the respective severity of liver disease/cirrhosis of the patient reported as CTP-score.

After establishing the CTP mapping the model was further validated via several comparisons with clinical data of ICG time courses in cirrhotic and control subjects (Fig. 6).

**Figure 6.**
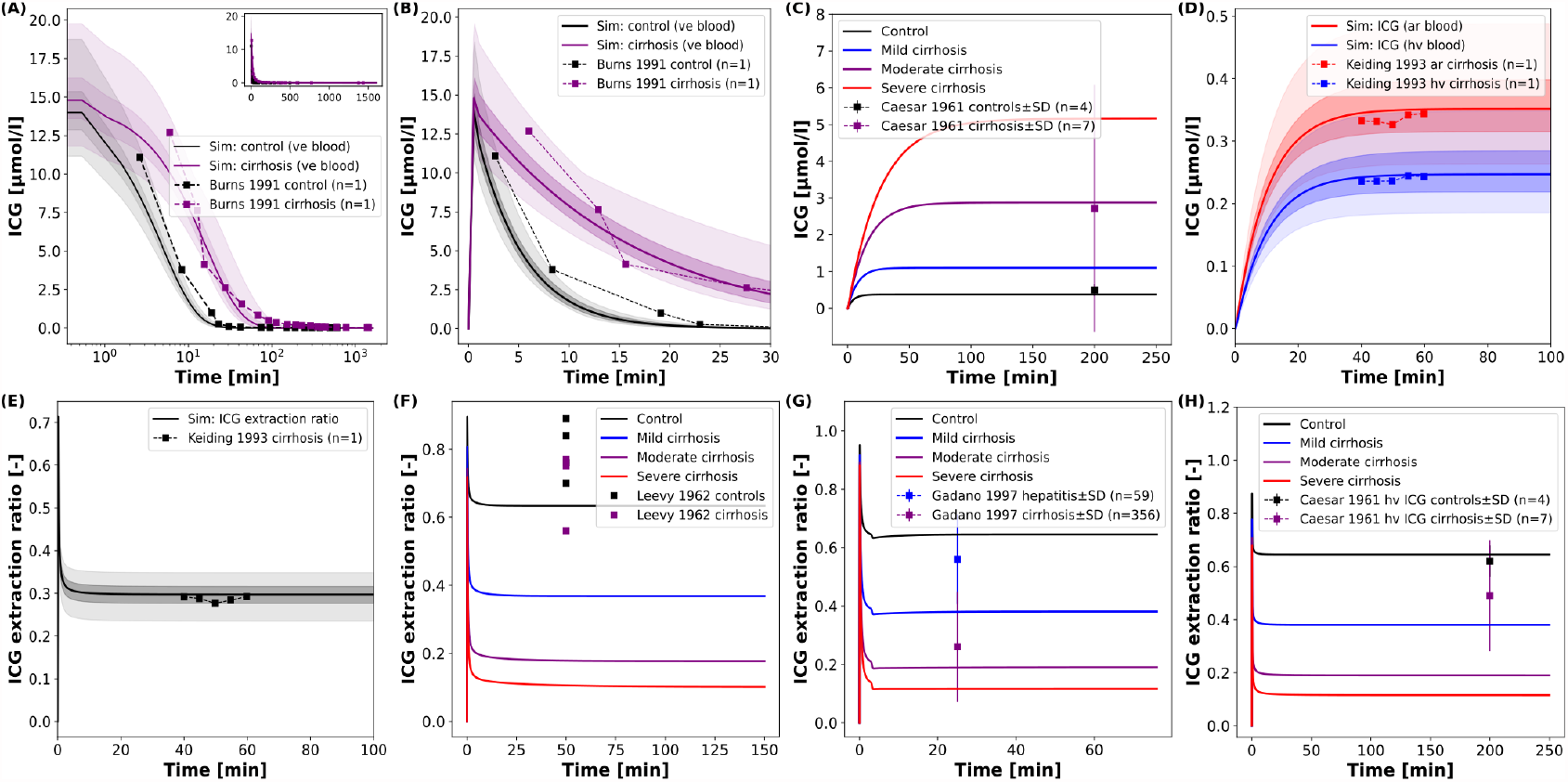
Model prediction of ICG time courses in subjects with cirrhosis: **A, B:** Venous concentration after a bolus ICG administration in a healthy subject and a cirrhotic patient (*f*_*cirrhosis*_ was set to 0.7 corresponding to moderate cirrhosis) (Burns et al., 1991). **C:** Venous concentration during a constant ICG infusion in healthy and cirrhotic subjects (Caesar et al., 1961). **D, E:** Hepatic venous and arterial ICG concentration and ICG extraction ratio in a cirrhotic patient (*f*_*cirrhosis*_ was set to 0.54 corresponding to mild-moderate cirrhosis) (Keiding et al., 1993). **F-H:** ICG extraction ratio in cirrhotic, hepatitis and control subjects during a constant ICG infusion (Leevy et al., 1962; Gadano et al., 1997; Caesar et al., 1961).

Assuming moderate cirrhosis (*f*_*cirrhosis*_ = 0.7), the model prediction of an ICG time course in a cirrhotic patient agrees well with the clinical data (Burns et al., 1991) (Fig. 6A,B). The main alteration compared to the healthy control is the slower disappearance rate resulting in higher ICG plasma concentrations. The same effect is observed in steady state via a constant ICG infusion (Fig. 6C). Using the *f*_*cirrhosis*_ values from the CTP mapping above, the steady state concentrations are predicted in agreement with the clinical data (Caesar et al., 1961). Fig. 6D,E shows the relation between the hepatic venous and arterial ICG concentrations and the extraction ratio in a cirrhotic subject. Here, *f*_*cirrhosis*_ was set to 0.54 which allowed to predict arterial and hepatic vein concentration as well as ICG extraction ratio. Finally, in Fig. 6F-H the ICG extraction ratio predicted for controls and three different cirrhosis degrees was compared to clinical data (Leevy et al., 1962; Gadano et al., 1997; Caesar et al., 1961). The extraction ratio in cirrhotic subjects is reduced compared to healthy controls, as predicted by the model.

### 3.5 Hepatectomy

After validating the model predictions of ICG pharmacokinetics in liver cirrhosis, the model was applied to liver surgery. To analyze the effect of partial hepatectomy on ICG elimination the change in ICG pharmacokinetic parameters as a function of the resection rate was simulated (Fig. 7A-D). The scan was performed for healthy controls as well as three different degrees of cirrhosis.

**Figure 7.**
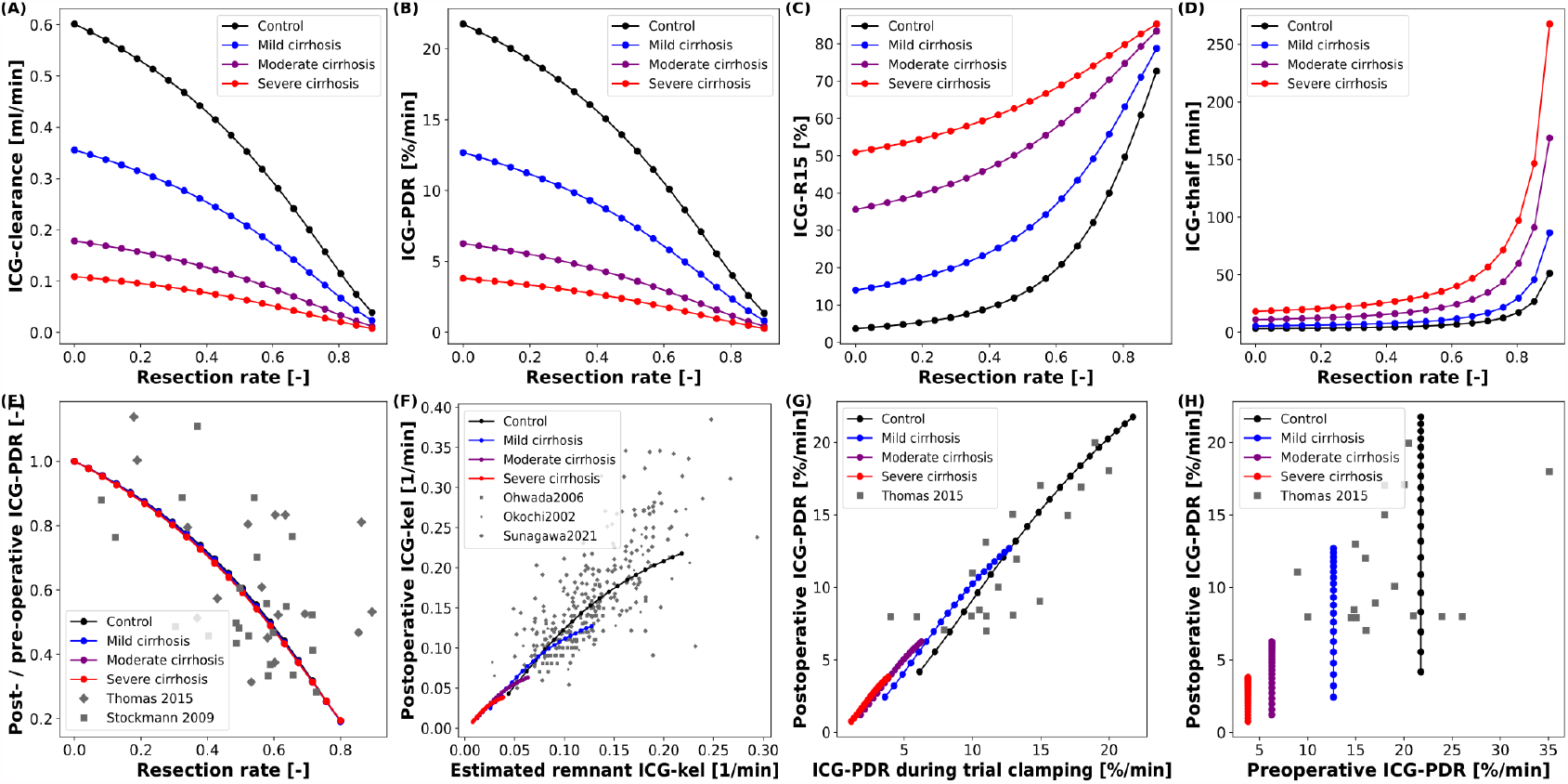
Model prediction of ICG pharmacokinetic parameters in hepatectomy under varying degree of cirrhosis: **A-D:** Dependency of postoperative ICG pharmacokinetic parameters on the resected volume in 4 different degrees of cirrhosis. **E:** Dependency of postoperative change in ICG-PDR on the resected volume (Thomas et al., 2015; Stockmann et al., 2009). **F:** Correlation between the measured postoperative ICG-kel and the estimated postoperative ICG-kel (product of preoperative ICG-kel and the future liver remnant) (Ohwada et al., 2006; Okochi et al., 2002; Sunagawa et al., 2021). **G:** Correlation between postoperative ICG-PDR and intraoperative ICG-PDR during trial clamping (Thomas et al., 2015). **H:** Correlation between postoperative and preoperative ICG-PDR (Thomas et al., 2015). Simulations for **E-H:** were performed for varying resection rates in healthy subjects and three different degrees of cirrhosis.

ICG-clearance and ICG-PDR are highest in the preoperative liver (resection rate = 0) and decrease with increasing resection rate whereas ICG-t_1/2_ and ICG-R15 are lowest in the healthy liver and increase with increasing resection rate. The effect of varying the degree of cirrhosis is in accordance with the results shown in (Fig. 4A-D). Importantly, increasing resection rate and increasing degree of cirrhosis affect ICG pharmacokinetic parameters in the same manner. The dependencies of ICG-clearance, ICG-PDR, ICG-t_1/2_ and ICG-R15 on the resection rate are fairly linear up to 50-60% resection, and become much more non-linear for higher resection rates.

For model validation the predictions were compared to clinical data of subjects undergoing hepatectomy. For these simulations the resection rate was varied from 0 to 0.9. First, the relative change of ICG-PDR after partial hepatectomy as a function of the resection rate was simulated (Fig. 7E). The model predicts a nonlinear dependency of change in ICG-PDR on the remnant liver volume independent of the degree of cirrhosis. This prediction is in good agreement with the clinical data (Thomas et al., 2015; Stockmann et al., 2009). Furthermore, the correlation between measured postoperative ICG-kel and estimated remnant ICG-kel ·(ICG-kel · fractional liver remnant) was simulated under various degrees of cirrhosis (Fig. 7F). A good correlation can be observed. The model predictions were compared to three different data sets (Ohwada et al., 2006; Okochi et al., 2002; Sunagawa et al., 2021) and are in good agreement with them. In addition, all data sets are in good agreement with each other. The simulated correlation line is independent of the cirrhosis degree, but with increasing cirrhosis ICG-kel decreases. A large variability can be observed in the experimental data, but as our simulations indicate is most likely not due to the underlying liver disease (cirrhosis).

(Thomas et al., 2015) found significant correlation between post-hepatectomy ICG-PDR and intraoperative ICG-PDR measured under trial clamping of those parts of the liver that were to be removed. This was simulated by changing hepatic blood flow and liver volume in separate simulations but in the same intervals. This was performed for a healthy liver as well as three different degrees of cirrhosis (Fig. 7G). The predictions agree well with the clinical data and show that reducing hepatic blood flow (clamping of liver volumes which will be resected) has a very similar effect on ICG elimination as actually removing the respective liver volume via hepatectomy.

Finally, the correlation between preoperative and postoperative ICG-PDR for different resection rates and cirrhosis degrees was simulated and compared to clinical data (Fig. 7H). ICG-PDR is reduced in cirrhosis preoperatively as well as postoperatively. The model prediction agrees with the clinical data (Thomas et al., 2015).

Overall the predictions of liver resection in severely cirrhotic liver is not in good agreement with the clinical data. This reflects the fact that no resections are performed in severely cirrhotic liver due to high risk of postoperative complications. As a consequence, most of the liver resection are performed in mild to moderate cirrhosis. The model allows to perform these risky hepatectomies *in silico* and predict there effect.

In summary, the model allows to systematically predict the changes of ICG pharmacokinetic parameters in HPB surgery under various degrees of liver disease (cirrhosis).

### 3.6 Prediction of post-hepatectomy survival

An interesting application of the presented PBPK model is the prediction of postoperative outcome for patients undergoing hepatectomy. Preoperative ICG-R15 and the planned resection rate are key parameters included in the decision process whether a patient is eligible to receive liver resection surgery.

As shown above, the presented PBPK model accurately predicts ICG-R15 in liver cirrhosis as well as the changes in ICG-R15 following hepatectomy. As such, we were interested how a classification model based on the PBPK model prediction of postoperative ICG-R15 compares to classification approaches only using clinical data (preoperative ICG-R15, resection rate and calculated postoperative ICG-R15).

Five alternative classification models were developed to predict survival after partial hepatectomy using a dataset of 141 patients (109 survivors and 32 non-survivors) (Seyama and Kokudo, 2009; Wakabayashi et al., 2004). Three of the classification models were solely based on clinical data: (i) Data1A - using preoperative ICG-R15, (ii) Data1B - using the calculated postoperative ICG-R15 by multiplying the future liver remnant (1-resection rate) with the preoperative ICG-R15; and (iii) Data2 - using both the resection rate and the preoperative ICG-R15. In addition two classification models were developed using PBPK model predictions as input: (iv) PBPK1 - using the predicted postoperative ICG-R15. Hereby, the model parameter *f*_*cirrhosis*_ was estimated for every subject based on preoperative ICG-R15, and postoperative ICG-R15 was predicted using *f*_*cirrhosis*_ and the corresponding resection rate; and (v) PBPK2 - using the resection rate and the estimated *f*_*cirrhosis*_ model parameter. An overview of the classification results of these five models is provided in Tab. 3, Fig. 8 and Fig. S1.

**Figure 8.**
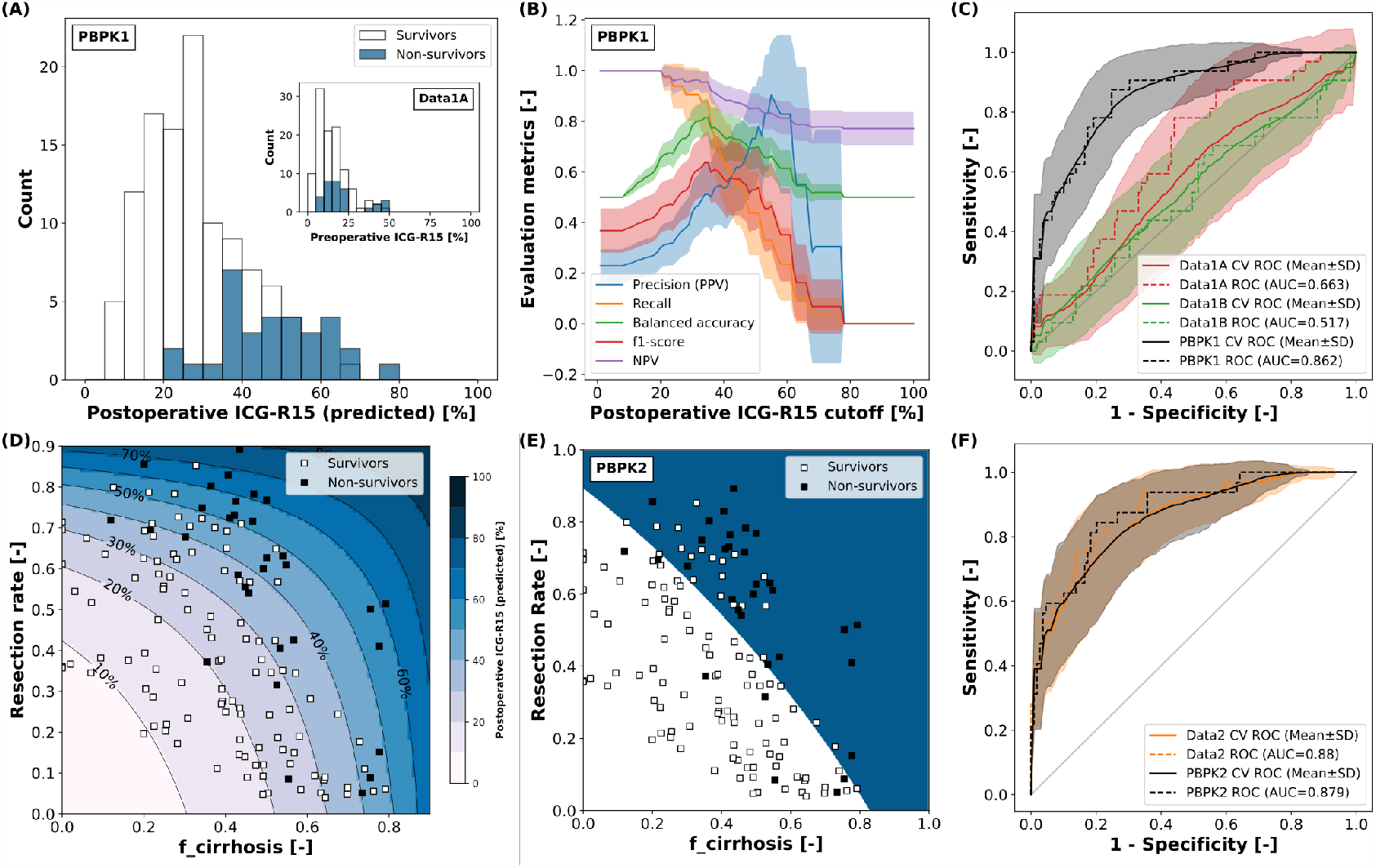
Classification of survivors/non-survivors after hepatectomy: All classification models predict non-survival after hepatectomy. Classification data (n=141) from (Seyama and Kokudo, 2009; Wakabayashi et al., 2004). **A:** PBPK model based prediction of postoperative ICG-R15 in survivors and non-survivors. *f*_*cirrhosis*_ was estimated for every subject based on preoperative ICG-R15 and post-operative ICG-R15 calculated using the respective resection rate (used in PBPK1). Corresponding measured preoperative ICG-R15 as inset (used in Data1A). **B:** Dependency of the evaluation metrics of the cross-validated classification model PBPK1 on the predicted postoperative ICG-R15 cutoff (mean ± SD). PPV: positive predictive value; NPV: negative predictive value **C:** ROC curve for the prediction of non-survival after hepatectomy using the complete dataset with cross-validation (mean ± SD) for classification models Data1A, Data1B and PBPK1. **D:** Predicted postoperative ICG-R15 and survival status depending on the resection rate and *f*_*cirrhosis*_ (used in PBPK2). *f*_*cirrhosis*_ was estimated for every subject individually based on preoperative ICG-R15 and post-operative ICG-R15 calculated using the respective resection rate. Shaded blue areas correspond to respective predicted postoperative ICG-R15. **E:** Decision boundary of the two-dimensional classification model PBPK2 based on the resection rate and *f*_*cirrhosis*_ using the complete dataset. White area: predicted survivor; blue area: predicted non-survivor. **F:** ROC curve for the prediction of non-survival after hepatectomy using the complete dataset with cross-validation (mean SD) for classification models Data2 and PBPK2.

Both PBPK-based classifiers (PBPK1, PBPK2) as well as the Data2 classifier outperform the data-based classifiers using a single feature (Data1A, Data1B) in predicting survival after partial hepatectomy.

When comparing the classification models using a single feature (Data1A, Data1B, PBPK1) the physiological-based predicted postoperative ICG-R15 (PBPK1) clearly outperforms the preoperative (Data1A) as well as calculated postoperative ICG-R15 (Data1B).

Fig. 8A shows the postoperative ICG-R15 in survivors and non-survivors predicted by the model as well as corresponding preoperative ICG-R15 (inset).

The PBPK model predicted postoperative ICG-R15 is able to distinguish better between survivors and non-survivors than the preoperative ICG-R15 as can be seen by the clearer separation of the histograms (Fig. 8A) and the respective ROC curves (Fig. 8C). Both preoperative ICG-R15 as well as calculated postoperative ICG-R15 are not very useful for the prediction of survival after partial hepatectomy, whereas predicted postoperative ICG-R15 using PBPK1 is a very good measure to predict survival after partial hepatectomy.

To determine possible cutoffs for predicted postoperative ICG-R15 based on the PBPK1 classifier the dependency of evaluation metrics on the cutoff was analyzed (Fig. 8B). Balanced accuracy as well as f1-score have a maximum at around 35%. The negative predictive value (NPV) as well as recall is 1 up to a predicted postoperative ICG-R15 ¡ 20%.

Fig. 8D depicts how the predicted postoperative ICG-R15 depends on the resection rate and *f*_*cirrhosis*_. The data confirms that a cutoff value slightly below 40% would correctly predict most of the non-survivors with a stricter cutoff of 20% avoiding any death after partial hepatectomy. Similar analysis of the data-based single feature classification models failed to find a significant optimum of evaluation metrics for either preoperative ICG-R15 or calculated postoperative ICG-R15 (see Supplemental Figure S1).

The two-feature classification models (PBPK2, Data2) show good performance in the survival prediction comparable to PBPK1 (Fig. 8E, F). Whereas the one-dimensional PBPK1 classifier provides a simple interpretation and cutoff value, the two dimensional classifiers are more difficult to interpret.

In summary, we developed a single-feature classification model based on a physiological-based model of ICG elimination (PBPK1) which allows to predict post-hepatectomy survival solely based on preoperative ICG-R15 input. Importantly, this computational model-based approach clearly outperforms data-based approaches such as preoperative ICG-R15 and calculated postoperative ICG-R15.

## 4 DISCUSSION

In summary, a PBPK model for ICG based liver function evaluation was developed, validated, and applied to the prediction of postoperative outcome after liver surgery, i.e., survival after partial hepatectomy. The model takes into account physiological factors such as the degree of cirrhosis and the estimated functional liver remnant, which allowed an accurate prediction of postoperative liver function in agreement with clinical data. As such, the model has proven its potential of becoming a valuable clinical tool for the planning hepato-pancreatico-biliary surgery.

The physiologically-based modeling approach allowed us to predict ICG pharmacokinetics data from 29 studies using only a small set of parameters and processes. The model accurately predicts changes in ICG pharmacokinetic parameters in a wide range of conditions including varying degrees of cirrhosis. Additionally, *in silico* hepatectomies with underlying cirrhosis are in good agreement with clinical data. As an important note, all clinical data besides the time courses in healthy subjects used for model calibration was used for model validation.

Overall, the clinical data shows large variability in ICG pharmacokinetic measurements, mostly due to inter-individual differences (e.g. Fig. 7F). Possible explanations are differences in blood flow, plasma proteins or protein amount or activity of the ICG transporter. An important next step would be a systematic analysis of these possible causes of variability and account for these confounding factors in the model.

One main outcome of this study is a single-feature classification model based on a physiological-based model of ICG elimination (PBPK1) which allows to predict post-hepatectomy survival solely based on preoperative ICG-R15 and resection rate. One limitation is the relatively small sample size (n=141) using retrospective data from Japanese subjects (Seyama and Kokudo, 2009; Wakabayashi et al., 2004). Further validation of our results with a dataset consisting of Caucasian subjects would be highly relevant.

The developed classification models demonstrated the potential of using PBPK predicted postoperative ICG-R15 values in the clinical decision process. Whereas the PBPK1 classifier provides a simple cutoff based on the individual model prediction (Fig. 8AB), PBPK2 provides the dependency of predicted postoperative ICG-R15 on resection rate and cirrhosis degree (Fig. 8D), both key factors for survival after liver resection.

Comparing different approaches of predicting postoperative outcome after partial hepatectomy showed the importance of taking resection rate into account. The data-based classifier combining resection rate with preoperative ICG (Data2) allowed to achieve comparable classification results to the PBPK-based classifiers. In contrast, the classification models Data1A and Data1B failed to achieve good prediction results.

Clinical data supporting our results have been reported by Haegele et al. (Haegele et al., 2016) who showed that an ICG-R15 *>*20% on postoperative day 1 predicted poor postoperative outcome, which agrees well with the results shown in Fig. 8B. The cutoff of ICG-R15 *<*20% allows to identify low-risk patients that are unlikely to have poor postoperative outcome after partial hepatectomy. This was confirmed by the high negative and low positive predictive values (¿80% and 30% respectively), suggesting that ICG-R15 is especially useful for the identification of low-risk patients. A recommendation was that subjects with ICG-R15 20-40% should undergo a more careful evaluation of the treatment options and additional information should be taken into consideration. In agreement with our results, Haegele et al. could show a significant improvement of the prediction of postoperative survival when using postoperative ICG-R15 (AUC_*roc*_=0.893) compared to preoperative ICG-R15 (AUC_*roc*_=0.719). As a side note, the experimental cutoff of *>*20% by Haegele et al. was determined in a Western population providing evidence that our classification approach could be generally applied despite developed on data from Japanese subjects.

Multiple studies showed only moderate performance in predicting post-hepatectomy outcome using preoperative ICG measurements. Gu et al. (Gu et al., 2020) found that preoperative ICG-R15 achieved an AUC_*roc*_ of 0.657 (95% CI 0.576-0.739) and 0.640 (95% CI 0.445-0.836) for the prediction of post-hepatectomy liver failure and 90-day mortality, respectively. Wong et al. (Wong et al., 2013) failed to achieve any significant prediction of postoperative severe morbidity using preoperative ICG-R15 (AUC_*roc*_=0.51, 95% CI 0.38-0.72). Wang et al. (Wang et al., 2018) found that preoperative ICG-R15 surpassed both CTP-score and MELD for the prediction of severe post-hepatectomy liver failure, but with moderate AUC_*roc*_=0.724 (95% CI 0.654-0.787). In summary, this data provides a strong argument for our approach of using predicted postoperative ICG-R15 values via a PBPK model for predicting postoperative survival, which allowed to improve the discriminatory power (see Fig. 8A).

Due to the high mortality rate, extended liver resection in the presence of cirrhosis is considered to be contraindicated. Recommendations are often that only selected patients with Child’s A status or preoperative ICG-R15 of less than 10% undergo major hepatectomy (Kitano and Kim, 1997). As can be seen in Fig. 8A (inset), even such a strict cutoff can still result in mortality after hepatectomy. We suggest using a combination of resection rate and individual liver damage (cirrhosis degree) estimated from preoperative ICG-R15 instead of relying on preoperative ICG-R15 alone. As shown by the PBPK2 classifier (and indirectly by the PBPK1 classifier) this approach allows a better evaluation of postoperative risk.

A very interesting result is that our model accurately predicted intraoperative ICG measurements (during trial clamping) (Fig. 7G), in which the blood flow to the area to be resected was clamped. These intraoperative ICG measurements are in very good agreement with post-operative ICG measurements (Thomas et al., 2015). Such an approach would allow to measure the expected postoperative ICG-R15 intraoperatively. Additional clamping data showed that the postoperative hospital stay was significantly longer for intraoperatively clamped ICG-R15 *>*20% (27.5± 14.1 days) compared to *<*20% (17.9*±* 9.2 days). Due to the good correlation between intraoperative clamped and postoperative ICG measurements this data provides additional support for the proposed lower cutoff of predicted postoperative ICG-R15 *<*20%. It is likely that postoperative ICG-R15 is not only a good predictor for survival but for postoperative complications in general as indicated by the data from Akita et al. (Akita et al., 2008). Additional support comes from Haegele et al. which showed significantly reduced liver dysfunction (3.6 vs 42.9%, p=0.001), reduced severe morbidity (16.1 vs 42.9%, p=0.016) and hospitalization (7 vs 11 days, p=0.019) for POD1 ICG-R15 *<*20% vs POD1 ICG-R15 *>*20% (Haegele et al., 2016).

The presented PBPK model predicts postoperative ICG-R15 immediately after partial hepatectomy, corresponding to clinical measurements on the first postoperative day (POD1). Using postoperative ICG-R15 on POD1 is supported by Haegele et al. (Haegele et al., 2016) who showed that an ICG clearance test was able to predict poor postoperative outcome as early as POD1.

Importantly, due to the physiologically based modeling approach our predictions could easily be further individualized with the availability of respective data. A personalized risk prediction based on an individualized PBPK model could include (i) general information such as age, sex and ethnicity; (ii) physiological information such as body weight, body fat percentage, cardiovascular parameters and organ volumes; (iii) information regarding the liver specifically such as liver perfusion, liver volume and quantification of ICG protein amounts as well as assessment of liver disease such as degree of cirrhosis would be of high relevance. Such an individualization could substantially improve the models prediction of postoperative liver function and outcome in patients undergoing partial hepatectomy. Going forward, an important next step will be to evaluate the model in the clinical context using a high quality dataset reporting individual ICG time courses in combination with a subset of the above-mentioned additional clinical data.

## AUTHOR CONTRIBUTIONS

AK and MK designed the study, developed the computational model, build the classifiers, implemented and performed the analysis, and wrote the initial draft of the manuscript. JG provided support with PK-DB and data curation. All authors discussed the results. All authors contributed to and revised the manuscript critically.

## DATA AVAILABILITY STATEMENT

The datasets generated for this study can be found in PK-DB available from https://pk-db.com.

