## Supplemental Figure 1 for "Prediction of survival after partial hepatectomy using a physiologically based pharmacokinetic model of indocyanine green liver function tests"

### Supplementary Material

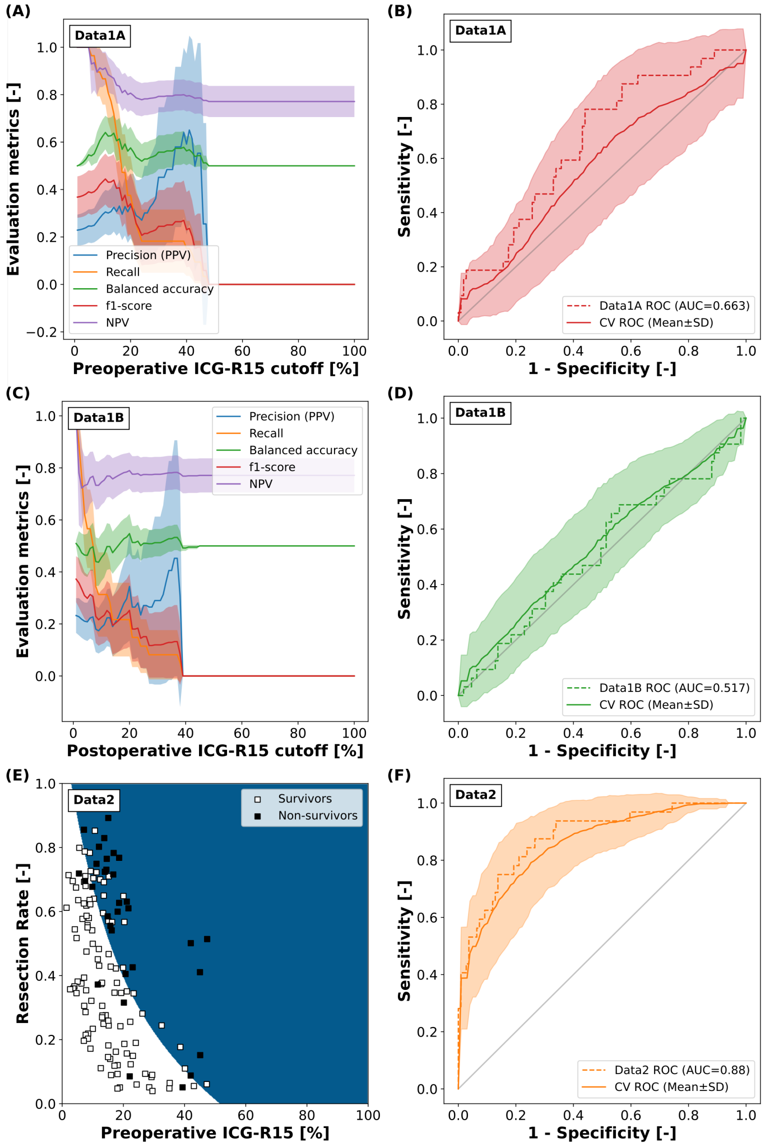


**Figure S1. Classification of survivors/non-survivors after hepatectomy**: **A**: Dependency of the evaluation metrics of the cross-validated classification model Data1A on the preoperative ICG-R15 cutoff (mean ± SD). PPV: positive predictive value; NPV: negative predictive value. **B:** ROC curve for the prediction of non-survival after hepatectomy using the complete dataset with cross-validation (mean ± SD) for classification model Data1A. ROC curve using the complete dataset (n=141) with cross-validation (mean ± SD) for classification model Data1A. **C**: Dependency of the evaluation metrics of the cross-validated classification model Data1B on the calculated postoperative ICG-R15 cutoff (mean ± SD). PPV: positive predictive value; NPV: negative predictive value. **D:** ROC curve for the prediction of non-survival after hepatectomy using the complete dataset with cross-validation (mean ± SD) for classification model Data1B. **E:** Decision boundary of the two-dimensional classification model Data2 based on the resection rate and $f_{cirrhosis}$ using the complete dataset. White area: predicted survivor; blue area: predicted non-survivor. **F:** ROC curve for the prediction of non-survival after hepatectomy using the complete dataset with cross-validation (mean ± SD) for classification model Data2.
